# The landscape of mouse epididymal cells defined by the single-cell RNA-Seq

**DOI:** 10.1101/2020.01.05.895052

**Authors:** Jianwu Shi, Mengmeng Sang, Gangcai Xie, Hao Chen

**Affiliations:** Institute of Reproductive Medicine, Medical School of Nantong University, Nantong, PR China

**Keywords:** Epididymis, scRNA-seq, Principal cell sub-populations

## Abstract

Spermatozoa acquire their fertilizing ability and forward motility properties during epididymal transit. Although lots of attempts elucidating the functions of different cell types in epididymis, the composition of epididymal tubal and cell types are still largely unknown. Using single-cell RNA sequence, we analyzed the cell constitutions and their gene expression profiles of adult epididymis derived from caput, corpus and cauda epididymis with a total of 12,597 cells. This allowed us to elucidate the full range of gene expression changes during epididymis and derive region-specific gene expression signatures along the epididymis. A total of 7 cell populations were identified with all known constituent cells of mouse epididymis, as well as two novel cell types. Our analyses revealed a segment to segment variation of the same cell type in the three different part of epididymis and generated a reference dataset of epididymal cell gene expression. Focused analyses uncovered nine subtypes of principal cell. Two subtypes of principal cell, c0.3 and c.6 respectively, in our results supported with previous finding that they mainly located in the caput of mouse epididymis and play important roles during sperm maturation. We also showed unique gene expression signatures of each cell population and key pathways that may concert epididymal epithelial cell-sperm interactions. Overall, our single-cell RNA seq datasets of epididymis provide a comprehensive potential cell types and information-rich resource for the studies of epididymal composition, epididymal microenvironment regulation by the specific cell type, or contraceptive development, as well as a gene expression roadmap to be emulated in efforts to achieve sperm maturation regulation in the epididymis.

## INTRODUCTION

The epididymis is a critical male sex organ that plays key roles in sperm transport, maturation and storage (Cooper and Yeung, 2010; Cornwall, 2009). Spermatozoa from testes acquires their motility and fertilization ability when they transit through the epididymis. In the epididymis, the sperm plasma membrane is subjected to sequential biochemical and proteomic modifications as a result of subtle interactions with components of the extracellular environment in the epididymal lumen.

The epididymis epithelium supports a luminal environment that promotes sperm maturation, and each region of the duct (caput, corpus, and cauda) is believed to play a unique role during sperm transition (Breton and Brown, 2013; Breton et al., 2016). The luminal secretions from caput and corpus epididymis are beneficial for the acquisition of sperm motility and fertilizing ability (Turner, 1995), cauda epididymal secretions are important for the proper storage of spermatozoa for several days in conditions that preserve their fertility (Jones and Murdoch, 1996). Each of these regions have been demonstrated to possess a distinct pattern of gene expression related to physiological functions which are important in the different steps of sperm maturation (Dube et al., 2007; Guyonnet et al., 2009; Johnston et al., 2005; Thimon et al., 2007). This compartmentalized gene expression triggers segment-specific protein secretions into the luminal fluid and, in turn, generates microenvironments optimized for each step of sperm maturation.

The composition of the intra luminal milieu is controlled by the surrounding pseudostratified epithelium, which is composed of five cell types, for now, possessing distinct physiological functions including principal cells, basal cells, clear/narrow cells, apical cells and halo cells all along the organ (Bernard Robaire, 2006; Cornwall, 2009; Sullivan et al., 2019). Studies of region-specific epididymal proteins showed that certain cell type was able to express quite different classification of genes, which contributes to the different physiologic functions of the segments (Cornwall, 2009; Dacheux and Dacheux, 2014). With the development of single-cell RNA sequencing (scRNA-seq), numbers of organs were analyzed in mammalian (Dong et al., 2018; Han et al., 2018; Tabula Muris et al., 2018), including male and female reproductive organ such as testis (Green et al., 2018; Guo et al., 2018; Wang et al., 2018) and ovary (Fan et al., 2019; Zhang et al., 2018) but not epididymis. However, the exact cell composition and gene expression repertoire of each cell population in epididymis are less characterized.

In this study, we applied microfluidic based scRNA-seq to the analyses of 12,597 cells derived from caput, corpus and cauda of mouse epididymis. Cell clustering analysis revealed five known cell types and more importantly discovered two novel epididymis cell sub-populations. Remarkably, nine sub-populations of principle cells were found in our study, which revealed unexpectedly heterogeneous in epididymis principle cells. Finally, our study represents the first regional transcriptome profiling of mouse epididymis at single cell level, which is important to the future study of the spatial microenvironments of the epididymis, and is also important for our understanding of epididymal diseases.

## MATERIALS AND METHODS

### Animals and epididymis sample collection

Sample collection was carried out under license in accordance with the Guidelines for Care and Use of Laboratory Animals of China and all protocols were approved by the Institutional Review Board of Nantong University. Five eight-week-old wildtype C57Bl/6J mice were used in this study. After sacrificing the mice, the epididymis were dissected and divided into three regions (caput, corpus, and cauda) and immediately stored in the GEXSCOPE^TM^ Tissue Preservation Solution (Singleron Biotechnologies) at 2-8 °C.

### Tissue dissociation and single cell suspension preparation

Prior to tissue dissociation, the specimens were washed with Hanks Balanced Salt Solution (HBSS) for three times and minced into 1–2 mm pieces. The tissue pieces were digested in 2ml GEXSCOPE^TM^ Tissue Dissociation Solution (Singleron Biotechnologies) at 37℃ for 15min in a 15ml centrifuge tube with continuous agitations. Following digestion, a 40-micron sterile strainer (Corning) was used to separate cells from cell debris and other impurities. The cells were centrifuged at 1000 rpm, 4°C, for 5 minutes and cell pellets were resuspended in 1ml PBS (HyClone). To remove red blood cells, 2 mL GEXSCOPE^TM^ Red Blood Cell Lysis Buffer (Singleron) was added to the cell suspension and incubated at 25°C for 10 minutes. The mixture was then centrifuged at 1000 rpm for 5 min and cell pellet re-suspended in PBS. Cells were counted with TC20 automated cell counter (Bio-Rad).

### Single cell RNA sequencing library preparation

The concentration of single-cell suspension was adjusted to 1×10^5^ cells/mL in PBS. Single cell suspension was then loaded onto a microfluidic chip (part of Singleron GEXSCOPE^TM^ Single Cell RNAseq Kit, Singleron Biotechnologies) and single cell RNA-seq libraries were constructed according to manufacturer’s instructions (Singleron Biotechnologies). The resulting single cell RNAseq libraries were sequenced on Illumina HiSeq X10 instrument with 150bp paired end reads.

### Single cell RNA-seq data analyses

In general, following steps were included in the analyses: raw sequencing data preprocessing, cell barcodes extraction, reads genomic alignment, unique molecular identifier (UMI) counting, cell sub-population discovery and sub-population gene markers finding. Firstly, reads with low sequencing qualities were filtered and the sequencing adapters were trimmed by Fastp (Chen et al., 2018) software (fastp 0.19.5) using default settings. Then, umi_tools (Smith et al., 2017) was used to identify and extract cell barcodes with the settings of cell number of 5000 and error correction threshold of 1. Next, STAR genomic mapper was applied to map the extracted reads to the mouse Gencode genome (GRCm38.primary_assembly.genome.fa, version M18). And then, featureCounts was used to assign exon-level reads based on Gencode gene annotation (gencode.vM18.primary_assembly.annotation.gtf, version M18). Furthermore, the UMIs for each gene were counted by umi_tools with the editing distance threshold of 1. Finally, cell sub-population discovery and sub-population markers finding were analyzed by Seurat 3.0 (Stuart et al., 2019), and only the cells with the number of feature RNAs between 500 and 6000, and the percentage of mitochondrial less than 0.5 were included for further analyses. The normalization of the expression data was performed by sctransform (SCT) algorithms (https://github.com/ChristophH/sctransform), and 3000 genes were selected for the integration of sub-populations from epididymis caput, corpus and cauda regions.

### Gene enrichment and transcription factor analyses

For each sub-population of epididymis or epididymal principle cells, the sub-population (or cell cluster) specific marker genes were selected for gene enrichment analyses, and the analyses were performed by clusterProfiler (Yu et al., 2012), which including gene ontology analysis. The file including 1675 mouse transcripton factors was downloaded from Riken Transcription Factor Database (http://genome.gsc.riken.jp/TFdb/). For each sub-population of epididymal principle cell, the average expression score was calculated based on SCT normalized expression values. The Maximal Regional Expression Difference (MRED) was defined as the difference between the maximal and the minimal expression scores among the three epididymis regions for all principle sub-populations. The 95% quantile value of the MREDs was used as the cutoff for the selection of the regional differentially expressed transcription factors.

## RESULTS

### ScRNA-seq analyses revealed a novel sperm interaction epididymal sub-population

Although quite a few studies elucidated the segment-specific function of epididymis (Belleannee et al., 2012; Sipila and Bjorkgren, 2016; Thimon et al., 2007), the comprehensive cell population of region-specific epididymal cells is still largely unknown. To define the cell types in mouse epididymis, we chose scRNA-seq to examine cells from fresh isolated caput, corpus and cauda region of the epididymis. A total of 12,597 cells from three regions of epididymis (Figure 1A and 1B) were conducted for scRNA-seq and subsequent analyses. Using Seurat analysis, we identified seven cell clusters with five known cell types in the mouse epididymis, including principal cell, clear/narrow cell, basal cell, sperm and T cells (Figure 1A and 1C, and more known marker gene expression in Figure S1).

**Figure 1.**
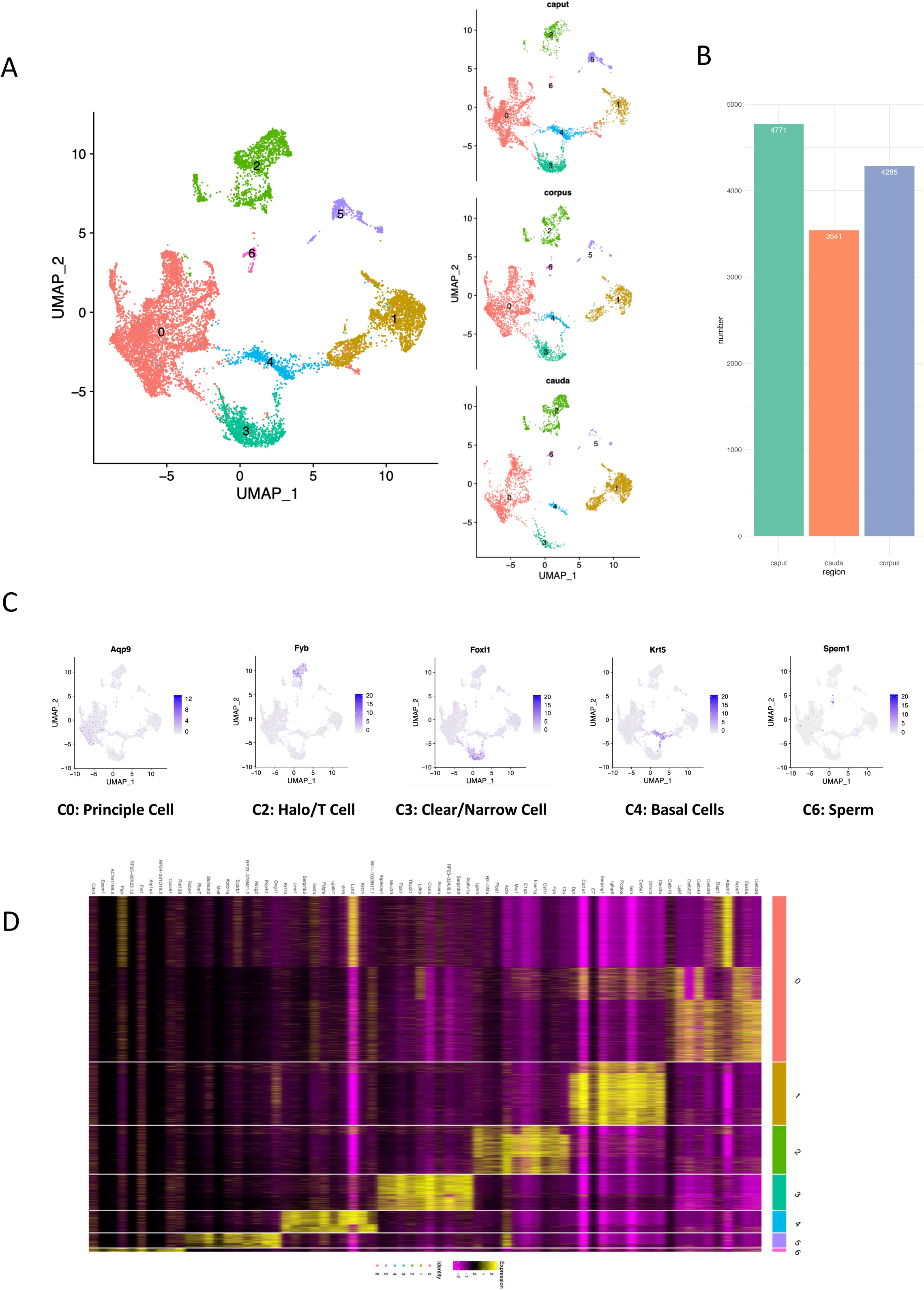
Single cell RNA sequencing of mouse epididymis regions. (A) Seven epididymis cell populations identified. Left: UMAP of the cell clustering based on all cells. Right: UMAP of the cell clustering based on cells from each epididymis region.(B) Number of single cells identified in each region.(C) Cell populations with known marker genes. (D) Top 10 marker genes for each cell population.

Regarding the cell clusters (C1 and C5) without known epididymis marker genes expressed, we did GO enrichment or DAVID gene enrichment analyses. Based on GO enrichment analysis, C5 was identified as vascular cells, the marker genes of which are enriched with “vascular development”, “regulation of angiogenesis” and so on (Figure S2). Interestingly, we identified a novel epididymis cell types (C1), the marker genes of which were enriched with DAVID terms “secreted”, “signal peptide”, “protein digestion and absorption” et al (Figure 2B). Furthermore, Epididymis Secretory Sperm Binding Protein (Col6a1) gene was highly expressed in this novel epididymis sub-population (C1), which indicates its possible interaction with sperm cells in epididymis (gene expression pattern illustrated in Figure 2A). And the highly specific expression of Olfactomedin Like 3 (Olfml3) in C1 novel cell type also revealed its possible intercellular functionalities (all top 10 C1 highly specific expressed genes can be found in Figure 2A). Interestingly, we found that the cell proportion of this novel epididymal sub-population (C1) was highest in epididymal cauda region, while lowest in caput region (Figure 2C).

**Figure 2:**
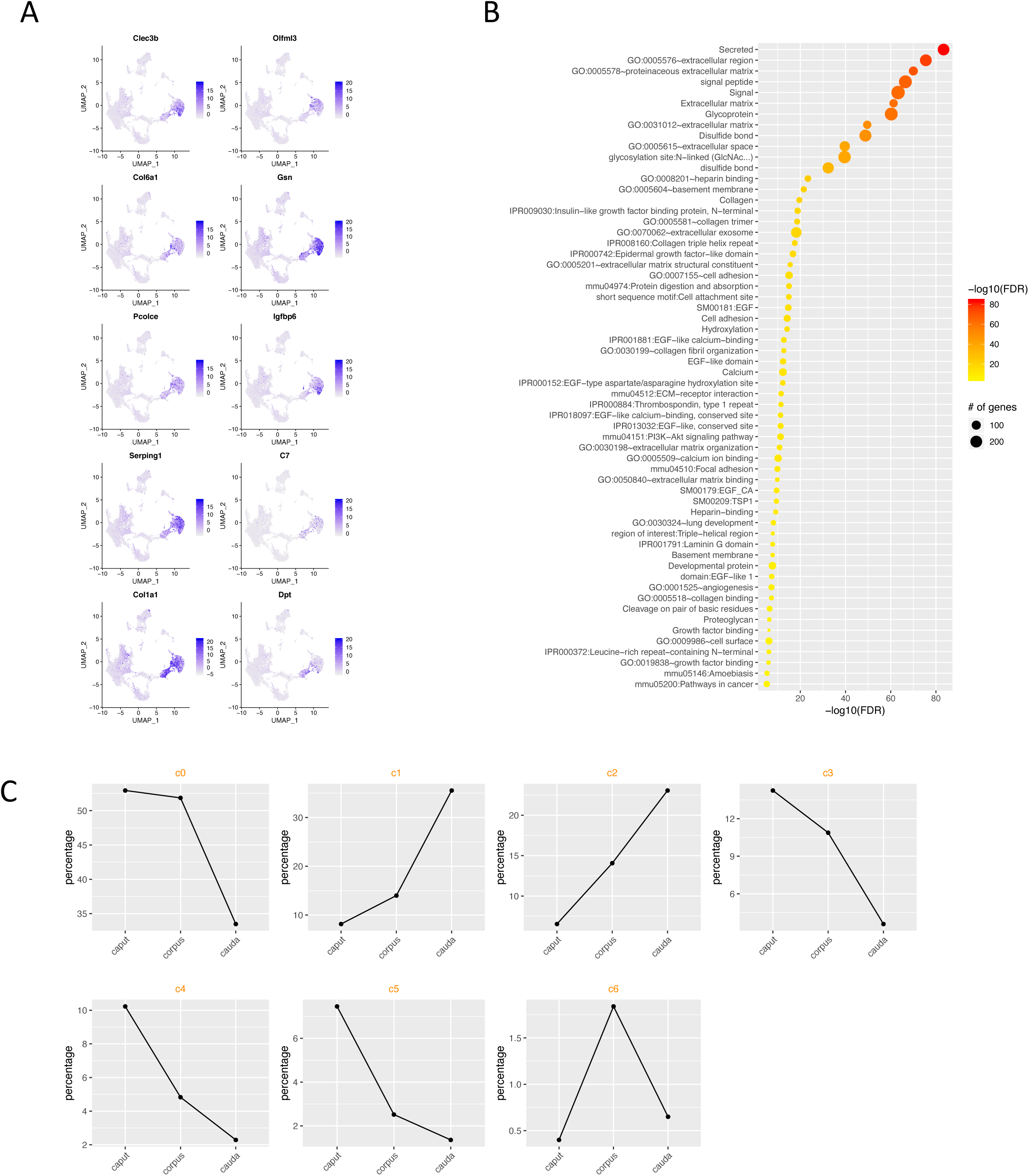
Study of the novel epididymis sub-population C1. (A) Specific expression of C1 sub-population marker genes. (B) Dotplot representation of DAVID enriched terms (DAVID terms selected based on FDR < 10^(-5)). (C) Increasing of C1 sub-population in the epididymal distal regions. The spatial proportion of each sub-population is also illustrated here.

For all the epididymis sub-populations (C0-C6), we identified population-level specifically expressed marker genes (Top 10 marker genes showed in Figure 1D, and all marker genes can be found in Table S1), which provides important marker genes for further study of epididymis sub-populations.

### Single cell RNA-Seq analyses recapitulate principal cell sub-populations

Our clustering analysis of the total epididymal cells suggested that the principal cell population is itself comprised of several sub-populations (Figure 1A). To further distinguish its sub-populations, we performed clustering on principal cells (Figure 3A). Nine different sub-populations of principal cell were revealed, which were defined as c0.1-c0.8 (Figure 3A). The pan-principal cell marker Aqp9 (Carvajal et al., 2018; Pastor-Soler et al., 2001) was found in all 9 clusters (Figure 3C), as expected. The enriched genes in each of these 9 principal cell were listed in the supplementary Table S2.

**Figure 3.**
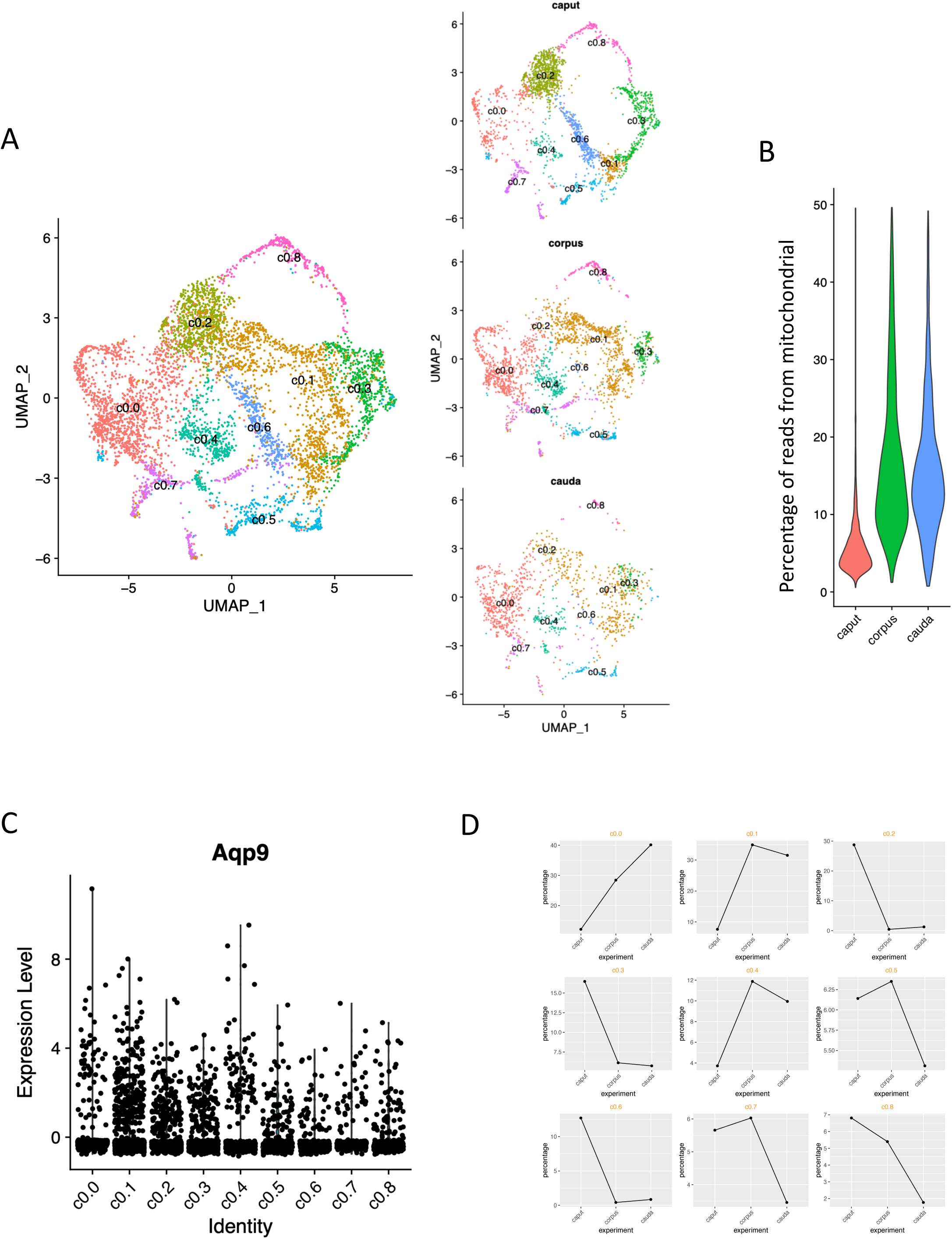
Sub-populations of principle cells identified. (A) UMAP representation of the sub-populations. Left: All single cells identified in principle cells (C0). Right: Split by epididymis regions. (B) Higher expression of mitochondrial genes in corpus and cauda. Only principle cells were included in this comparison. (C) Principle cell marker Aqp9 expression in each sub-population. (D) Patterns Of epididymis sub-population spatial distributions.

Principal cell is the major epididymal cell which composes the entire tubules and Aqp9 is the common recognized principal cell marker gene(Pastor-Soler et al., 2010). However, a comprehensive census of principal cell in the specific epididymal region has been hampered by their marker gene rarity. To address these issues, we applied single cell genetic strategies and screened out top ten marker genes for each subsets of principal cell (Figure 4A). Representative marker genes for each principal cell subsets were visualized in UMAP space (Figure 4B). Consistently, the cell population c0.3 and c0.6 have been reported in the previous studies (Ma et al., 2013; Xie et al., 2013), which were identified through the marker genes Lcn5 and Spink13 respectively (Figure 4A&B). Furthermore, the cell subset c0.3 was the most abundant cells at the caput region of epididymis with the high expressions of Defb 15, 29 and 30 (figure 5A&B). GO analysis also revealed its functional role of defense response, suggesting its predominant role in defensin family regulation (Figure 5C).

**Figure 4.**
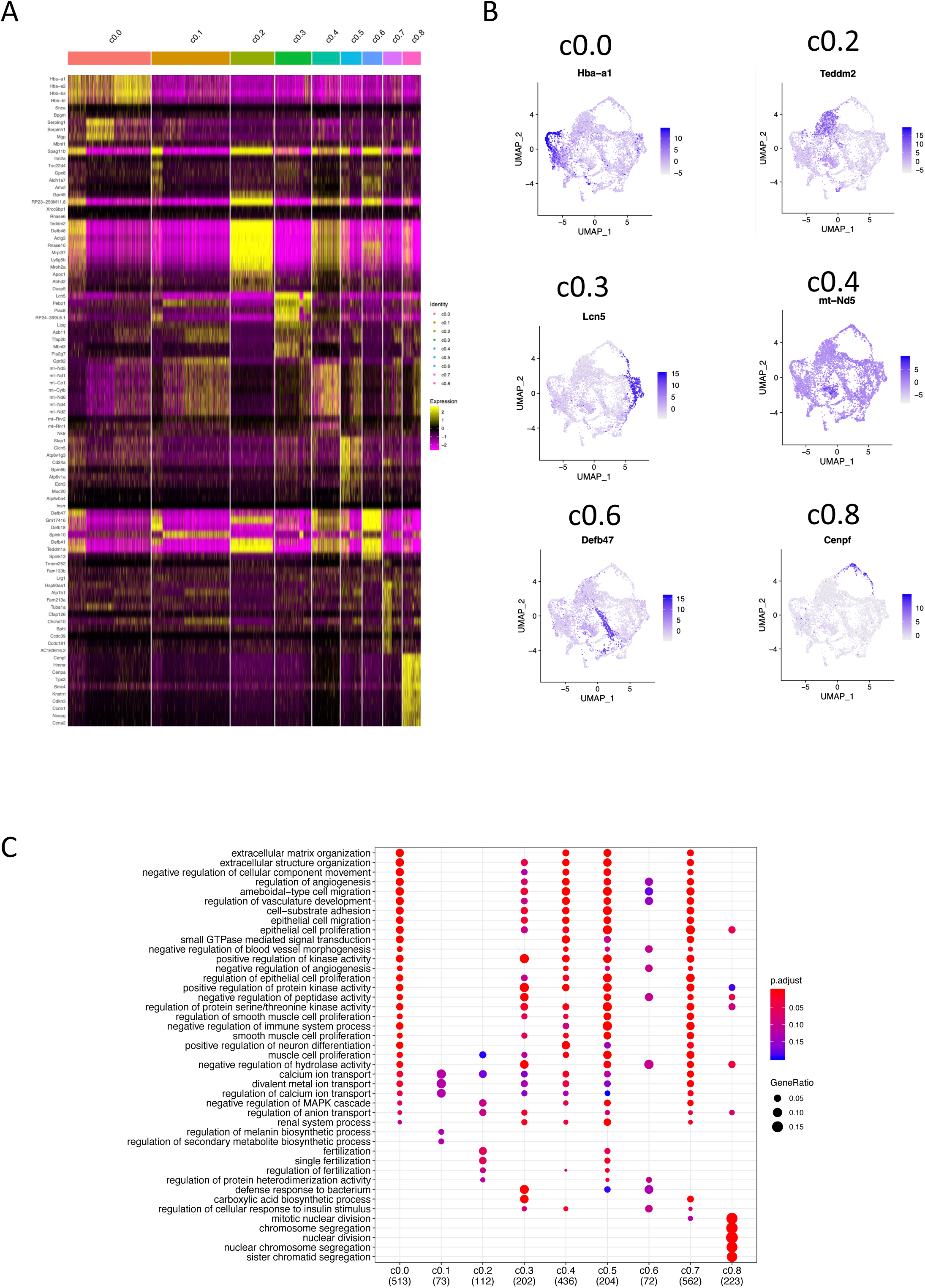
Features of principle cell sub-populations. (A) Heatmap showing top 10 marker genes for each sub-population. (B) UMAP showing representative marker genes for the principle sub-populations. (C) GO enrichment analysis for each sub-population.

**Figure 5.**
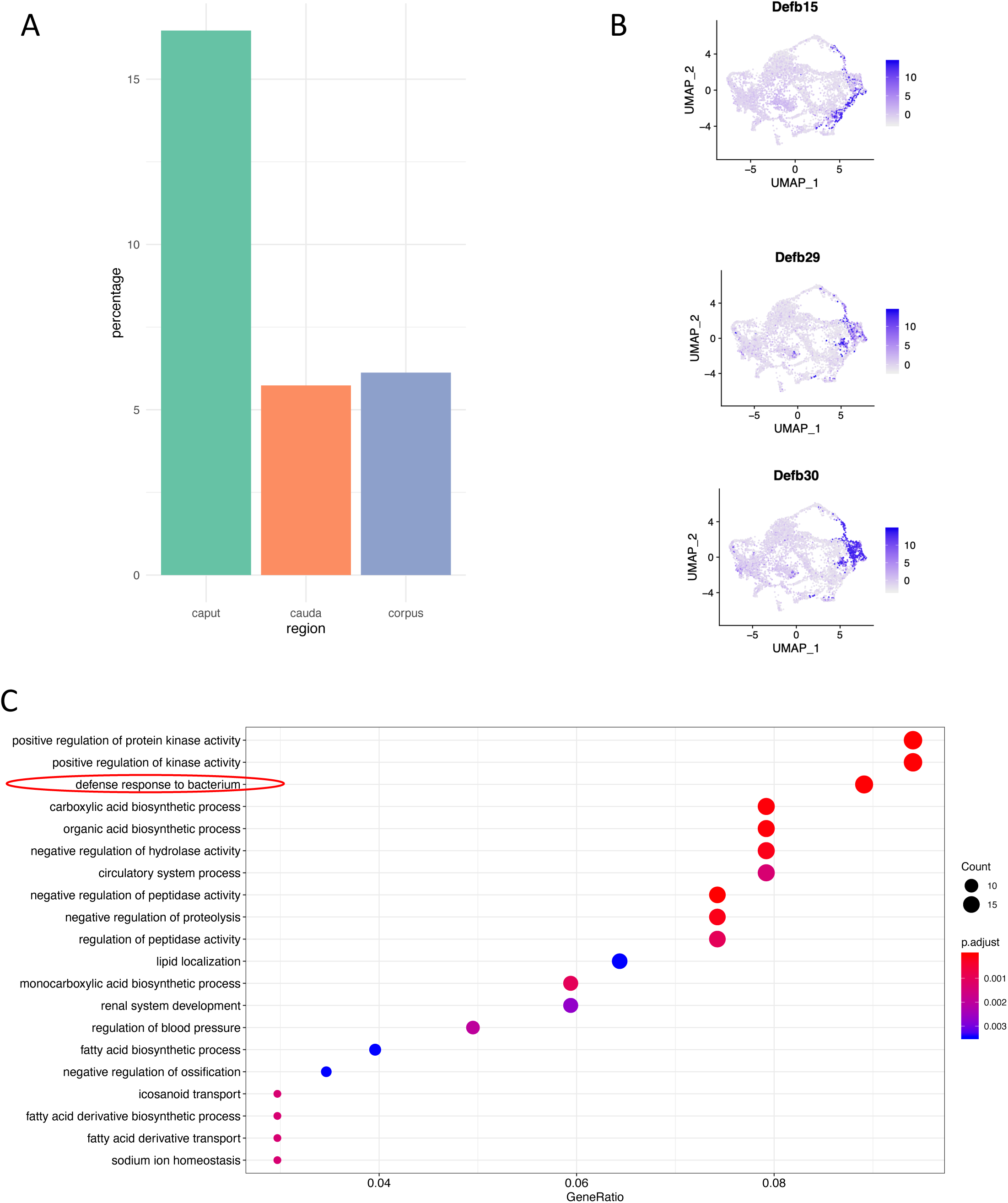
Study of principle cell c0.3 sub-population. (A) c0.3 epididymis sub-population is enriched in the caput region of epididymis. Y-axis: proportion of cells from c0.3 (total: cells in principle cell population) in each region. (B) Specific expression of Defensin genes in c0.3 principle sub-population. (C) GO enrichment analysis for the marker genes in c0.3 principle sub-population. Genes related to “Defense response to bacterium” are significantly enriched.

Apart from reported cell subsets, the seven remaining clusters represent novel populations, corresponding to extracellular structure organization(c0.0), small GTPase mediated signal transduction (c0.4), regulation of protein serine/threonine kinase activity(c0.5), epithelial cell proliferation (c0.7) and nuclear division or organelle fission(c0.8) process analyzed by GO enrichment. The represented GO terms associated with the differentially expressed genes in each cell cluster (Figure 4C) manifested diverse roles of the principal cell in regulating epididymal microenvironments, albeit three cell subsets remained unknown GO processes.

### Epididymal cell population proximal-distal distributions

In the total epididymal cell study, we found spatial distribution patterns of the seven epididymal cell populations. Four cell populations (C0: Principle Cell; C3: Clear/Narrow Cells; C4: Basal Cells; C5: Sperm cells) were distally reduced, while two populations (C1: Novel epididymal cell identified in this study, C2: Halo/T Cells) were distally increased (Figure 2C). Intriguingly, expressions of lots genes showed segment-specific or distally changed pattern (Bjorkgren and Sipila, 2019; Sipila and Bjorkgren, 2016), suggesting their expression patterns might be altered along with the cell number changes. The spatial distribution changes of each epididymal cell population might help to reveal their functionalities.

In the principle sub-population study, unexpectedly, we discovered that expression of mitochondrial genes was significantly higher in the epididymal corpus and cauda regions comparing to the caput region (Figure 3B). This observation suggests that the energy requirements in the distal epididymal regions might be stronger. For each principle sub-population, we also did cell population regional distribution studies. In total, three epididymal principle sub-populations (c0.0, c0.1, c0.4) were observed as distally increased, and six sub-populations (c0.2, c0.3, c0.5, c0.6, c0.7, c0.8) were distally decreased (Figure 3D).

Based on the total cell population and principle sub-population spatial distribution studies, we suggested that the different microenvironments of the three epididymal regions might be due to the cell population differences. And the epididymal proximal-distal microenvironments differences might also suggest region-dependent functionalities in mouse epididymis for sperm maturation.

### Regional regulation patterns identified in the principle sub-populations

Finally, we systematically studied the transcriptional regulations in mouse epididymal principle-cell sub-populations (Figure 6). Based on the global distribution of the MREDs for the expressed genes (Figure S3), we found 459 transcription factors (TFs) were spatially differentially expressed, and the expression of 177 TFs of them was distally decreased or increased (Table S3). We found genes, such as Enpp2 and Bmyc, were highly regional differentially expressed (Top 10 reginal differentially expressed TFs shown in Figure 6A). Enpp2 had been annotated as both of phosphodiesterase and phospholipase, and might be important for the catalyzing of the production of lysophosphatidic acid in extracellular fluids (ENPP2 in GENECARDS). Based on TF-TF expression correlation and network study (Figure 6B, F), Nupr1 gene was found to be highly correlated with other TFs, which indicates its important role in the principle cell regulation network. Interestingly, we found three patterns of regional expression of TFs, some TFs were both regionally and sub-population specifically expressed, such as Nupr1 and Pdlim1 (Figure 6C), some TFs such as Lhx1, Stat5b and Meis1 were highly specifically expressed at certain epididymal region and principle sub-population (Figure 6D), and some TFs such as Runx2, Scand1 and Mafb (Figure 6E) were highly reginal expressed no matter which principle sub-population they were in. This information highlighted the complex transcriptional regulation patterns in epididymal principle sub-populations, and the regulation of which dependents on both of the epididymal location and sub-population types.

**Figure 6.**
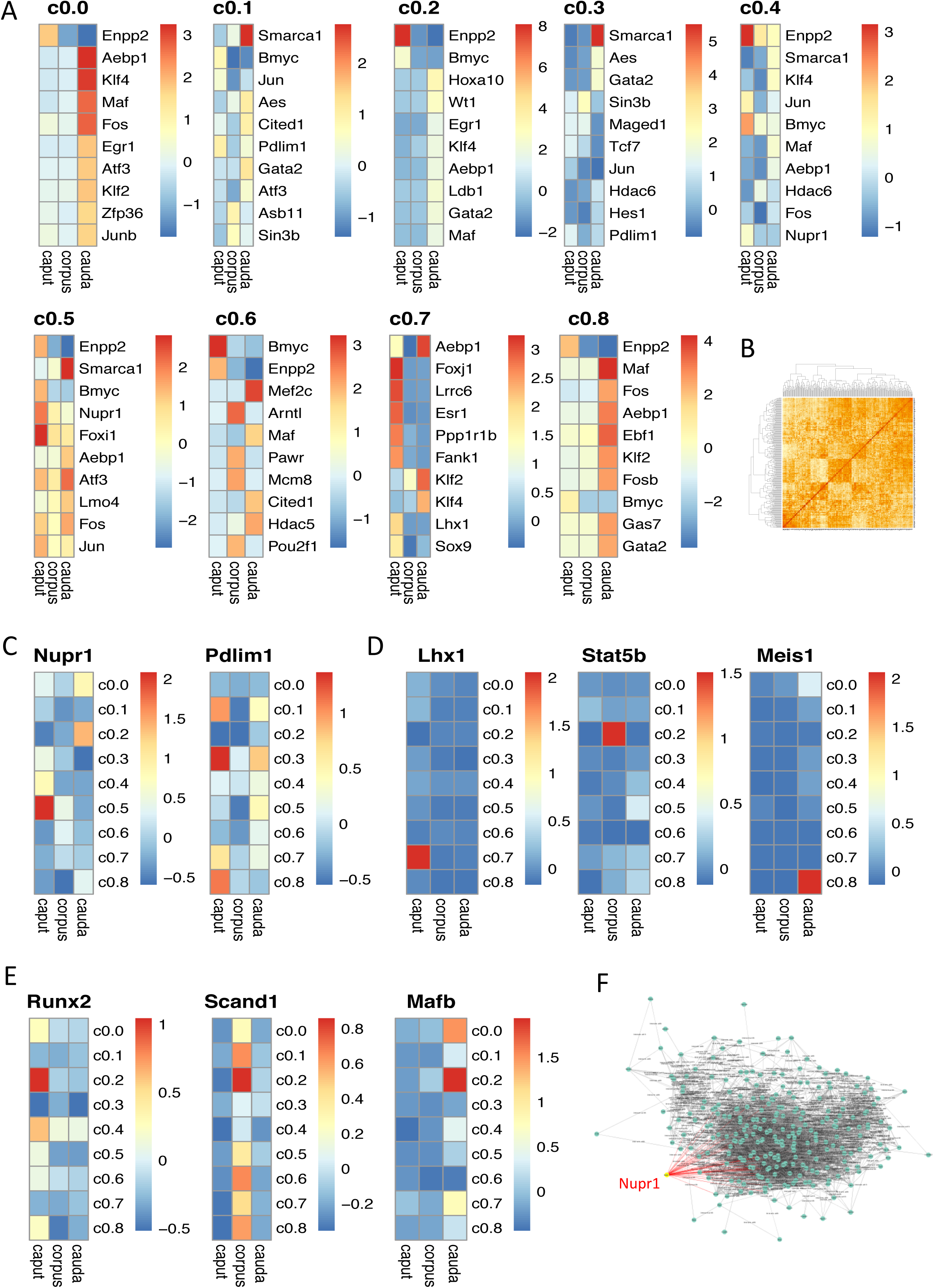
Regional regulation patterns in epididymal principle sub-populations. (A) Top 10 regional differentially expressed transcription factors (TFs) for each principle sub-population. (B) Heatmap representation of the correlation matrix for the regional differentially expressed TFs. Spearman’s *rho* was used for correlation calculation. (C) Examples of TFs with high regional and sub-population expression variations. (D) Examples of TFs with high specificities of regional and sub-population expression. (E) Examples of TFs specifically expressed in one epididymal region. (F) TF-TF correlation network showing the network of Nupr1.

## Discussion

The mammalian epididymis is composed of convoluted and interconnected regions, each of which contains a complex mosaic of spatially intermixed cells. Overwhelming studies have attempted to characterize the cell types in the epididymis based on cell location, morphology, connection, function, and marker gene expression (Da Silva et al., 2011; Shum et al., 2011; Shum et al., 2014). However, there is still a lack of knowledge regarding the epididymal cell compositions and its gene regulations from caput to cauda of epididymis. So we, for the first time, profiled the single cell RNA sequencing of the whole mouse epididymis, which included 12,597 epididymal single cells from caput, corpus and cauda regions. In total, we identified six epididymal cell populations according to known marker genes or GO enrichment analysis. And interestingly, we discovered one novel epididymal cell population that cannot be identified as known cell types. This novel cell population is enriched with Epididymis Secretory Sperm Binding Protein (Col6a1) and Olfactomedin Like 3 (Olfml3), which indicates its possible interaction with sperms and might play an important role in sperm maturation. Unexpectedly, the epididymal principal cell, as the major cell type in epididymis, can be further categorized into nine sub-populations, and all of which express classical principal marker gene Aqp9 (Pastor-Soler et al., 2010). Notably, those principal sub-populations exhibit proximal-distal distribution variations in mouse epididymis, which suggests their functional divergence for sperm maturation.

In the past decades, emerging studies endeavored to elucidate the regulation of region-specific gene expression in the epididymis and its potential contribution for sperm maturation using varies strategy including genetic regulation (Johnston et al., 2005; Oh et al., 2006; Zhang et al., 2006), small RNA(Anderson et al., 2015), epigenetic control(Sipila and Bjorkgren, 2016) and androgen-responsive regulations (Belleannee et al., 2012; Pihlajamaa et al., 2014). Region specific gene expression of epididymis in the mouse and human was also studied by microarray analysis(Johnston et al., 2005; Zhang et al., 2006). On the other hand, it was plausible that the epididymosome was contributed to the sperm modifications during the sperm transient in epididymis (Sharma et al., 2018; Trigg et al., 2019). Recently, James et al. demonstrated the expression profiles of segment-specific human epididymal epithelial cells (Browne et al., 2016). The human epididymal epithelial cells were isolated and differentiated cultured from caput, corpus and cauda and then underwent RNA-seq. However, the specific cell constitution and contribution which locate in the different region are less characterized. Our results depicted the cell landscape from each region of the adult mouse epididymis and in-depth analysis of the gene expression pattern of each cell population, suggesting more complexity roles of cell-specific regulation of epididymis besides segment-specific modulation.

Based on the current knowledge, the epididymal epithelium is composed of five distinct cell types: principal, narrow/clear, apical, Halo (T cell) and basal cells (Bernard Robaire, 2006; Cornwall, 2009; Sullivan et al., 2019). Supporting previous studies, we found all known cell types based on the reported cell-specific marker genes (Figure 1). In detail, Cell population C0 was defined as principal cell; cell population C2 was referred to as Halo cell; cell population C4 was corresponding to basal cell, as well as cell population C6 was identified as sperm. Cell population C3 was identified as Clear/Narrow cell due to lack specific cell markers. Intriguingly, the cell number of these known cell populations was divergent along with the different region. For example, cell population of C0, C3, C4 and C5 was gradually decreased, whereas cell population of C1 and C2 was elevated from caput to cauda. Convincingly, our results of cell population C3 dramatically reduced in corpus and cauda were in line with previous study that cell/Narrow cell were exclusively in the caput region (Bernard Robaire, 2006; Breton et al., 2016).

Besides specific expression of Col6a1 and Olfml3, the novel epididymal cell population C1 is also significantly enriched with extracellular structure organization, extracellular matrix organization and epithelial cell proliferation. The top 10 expressed genes in C1 are Clec3b, Olfml3, Col6a1, Gsn, Pcolce, Igfbp6, Serping1, C7 and Col1a1. Consisting with GO analysis, all these genes were predominantly localized in the extracellular compartment according to the information from GeneCards (Stelzer G, 2016). In particularly, C-Type Lectin Domain Family 3 Member B (clec3b) was reported to control cell proliferation in clear cell renal cell carcinoma(Liu et al., 2018) as well as low clec3b in exosomes derived from HCC promoted cell migration, invasion and epithelial–mesenchymal transition(Dai et al., 2019), indicating its contribution to the maintenance of epididymal epithelium. Recent study found that Gelsolin (Gsn) was the major secreted protein in the distal region of the bovine epididymis(Belleannee et al., 2011; Dacheux and Dacheux, 2014). In line with previous finding, our results demonstrated that the number of C1 cells was elevated from caput to cauda epididymis, suggesting its potential involvement for the sperm storage. According to the GO enrichment analysis, the population of C5 cells was correspond to the regulation of vasculature development and angiogenesis, which might be responsible for the communication between epididymal cells and vasculature cells the epididymis.

Specifically, principal cell is the most abundant cell type in the epididymal tubule, which accounts for ∼80% of the epithelium(Cornwall, 2009). However, little is known about the differences of region-specific principal cell as well as the transcriptome profiling in the three regions of epididymis. Our data demonstrated for the first time the multiple cell subsets of principal cells with distinct marker genes in each sub-population. Clustering analysis focusing on the principal cells derived from caput, corpus and cauda epididymis revealed nine major cell types, two of which had been reported to be located at the caput region, and such localization is also confirmed in our study. The marker gene Lcn5 was highly and specifically expressed in the sub-population of principal cells c0.3, which mainly located at the caput region of epididymis (Xie et al., 2013). And another marker gene spink13 in c0.6 cell population was expressed in the principal cell of initial segment of caput and was involved in the regulation of sperm acrosome reaction (Ma et al., 2013).

The traditional perspective of epididymis being a well differentiated organ is not thought to have stem cell, nor its epididymal epithelial cells to divide in adult (Bernard Robaire, 2006). The epididymal epithelial cells consists of four cell types including principal cells, basal cell, clear cells and narrow cells. Recently, several studies have been investigated the possible proliferation of epididymal epithelial cells both in vitro and in mammalian animals. The isolated bovine epididymal cells were stimulated to be proliferation when co-cultured with spermatozoa(Reyes-Moreno et al., 2008). Bernal-Manas et al. reported the cell proliferation of epididymal epithelium (mainly clear and principal cells) of *sus domesticus* epididymis with PCNA immunochemistry staining (Bernal-Manas et al., 2014). Another study demonstrated that the basal cells gained the capacity of proliferation after efferent duct ligation (EDL) in mice (Kim et al., 2015). The most recent studies found that the epididymal epithelia cells was able to rapidly grow after efferent duct ligation or orchidectomy. Basal cells was considered as a competitive candidate for the epididymal stem cell, albeit more convincing results need to be provided (Pinel et al., 2019). These results suggested the proliferative possibility of epididymal epithelial cells during epididymal cells dynamics. Of note, our data showed that cell subset c0.7 of principal cell was predicted as progenitor/stem cells with significantly enrichment in the epithelial cell proliferation and cell growth according to gene ontology analysis, suggesting that it might serve as stem cell or progenitor cells to keep the intact of epididymis. The proliferative cell population c0.7 from our data is comparable to previous results, although the cell source was fundamentally different. Future studies will aim to investigate whether the population c0.7 play a critical role in epididymal epithelium homeostasis and its potential stem cell capability.

Previous study reported the apical mitochondria-rich cells in both mouse and human epididymis These cells were believed to be distinct from principal cells and involved in the holocrine secretion and acidification of epididymal fluid (Martinez-Garcia et al., 1995). In contrast with previous studies, we surprisingly found the mitochondria percentage of principal cell was significantly increased in corpus and cauda region compared to caput region of epididymis. Furthermore, one of sub-population of principal cell (c0.4) is highly enriched with the expression of mitochondria genes such as mt-Nd1, mt-Co1, mt-Nd5, mt-Nd2, mt-Cytb, mt-Nd6, mt-Nd4, mt-Rnr2, Nktr and mt-Rnr1, most of which are believed to belong to the core subunits of mitochondrial membrane respiratory chain NADH dehydrogenase (Complex I) and Cytochrome C Oxidase complex, that is essential for the maintenance of mitochondrial functions and oxidative phosphorylation. Our data suggested the apical mitochondria-rich principle cell might play more important role in the corpus/cauda than caput region of epididymis. The detailed characterizations for the c0.4 cell sub-population still need to be further elucidated.

Epididymal cells secret plenty of the proteins, small RNA and extracellular vesicles into the epididymal lumen to regulate sperm maturation (Bjorkgren and Sipila, 2019). DNA binding transcription factors (TFs) play center roles in the regulation of gene expression. Therefore, we further explored the differentially expressed TFs and their expression profiles. Among TFs that were abundant in the majority of principal sub-population cells, some of them are related to epididymis function in our data, such as bmyc and enpp2. The bmyc, member of the Myc family to inhibit myc, was found to be predominately expressed in the caput region (Cornwall et al., 2001) and regulated by androgen and testicular factors. Bmyc gene knockout (KO) experiment showed that the KO mice had both of smaller testes and epididymis(Turunen et al., 2012). Enpp2 was detected in the epithelial cells in rat epididymis (Belleannee et al., 2010). Besides, recent study demonstrated the expression patterns of primary cultured human epididymal epithelial cells derived from caput, corpus and cauda regions (Browne et al., 2016). They reported a total of 90 deferentially expressed TFs associated with epididymis epithelial cells. Compatible with previous study, expression profile of several TFs was identical such as creb3l1, tbx3, atf3 etc. However, more TFs (eg. Hlf, esr1, pou2f2, fosb and cdkn2c) displayed the inconsistent with previous results. Reasons for the inconsistency might be on account of the different of species between human and mouse. On the other hand, cultured cells may not fully epitomize the cells in the original organ because of its loss of cell communications and other hormonal regulations.

Taken together, our data provide an indispensable resource, at single cell resolution, for a comprehensive cell atlas of the epididymis. Combine with further studies on the detailed mechanisms of epididymal cells communications and microenvironments hemostasis for sperm maturation, it will shed light on the understanding of normal epididymis functionalities and the cellular mechanisms of epididymis diseases that cause male infertility.

## Authors’ contributions

HC conceived and designed the experiments. JWS, MMS and GCX performed the experiments and analyzed the data. JWS, GCX and HC wrote the paper.

---

**Figure S1. Expression of known marker genes.** Marker genes from six known epididymis cell types were illustrated in UMAP plot.

**Figure S2. GO enrichment analysis for epididymis C5 cell population.** GO terms related to vascular were enriched.

**Figure S3. Density plot of MREDs of all expressed genes.** 95% quantile value is highlighted by green dash line.

