## Supplementary figures and images for "The landscape of mouse epididymal cells defined by the single-cell RNA-Seq"

### Figure S1

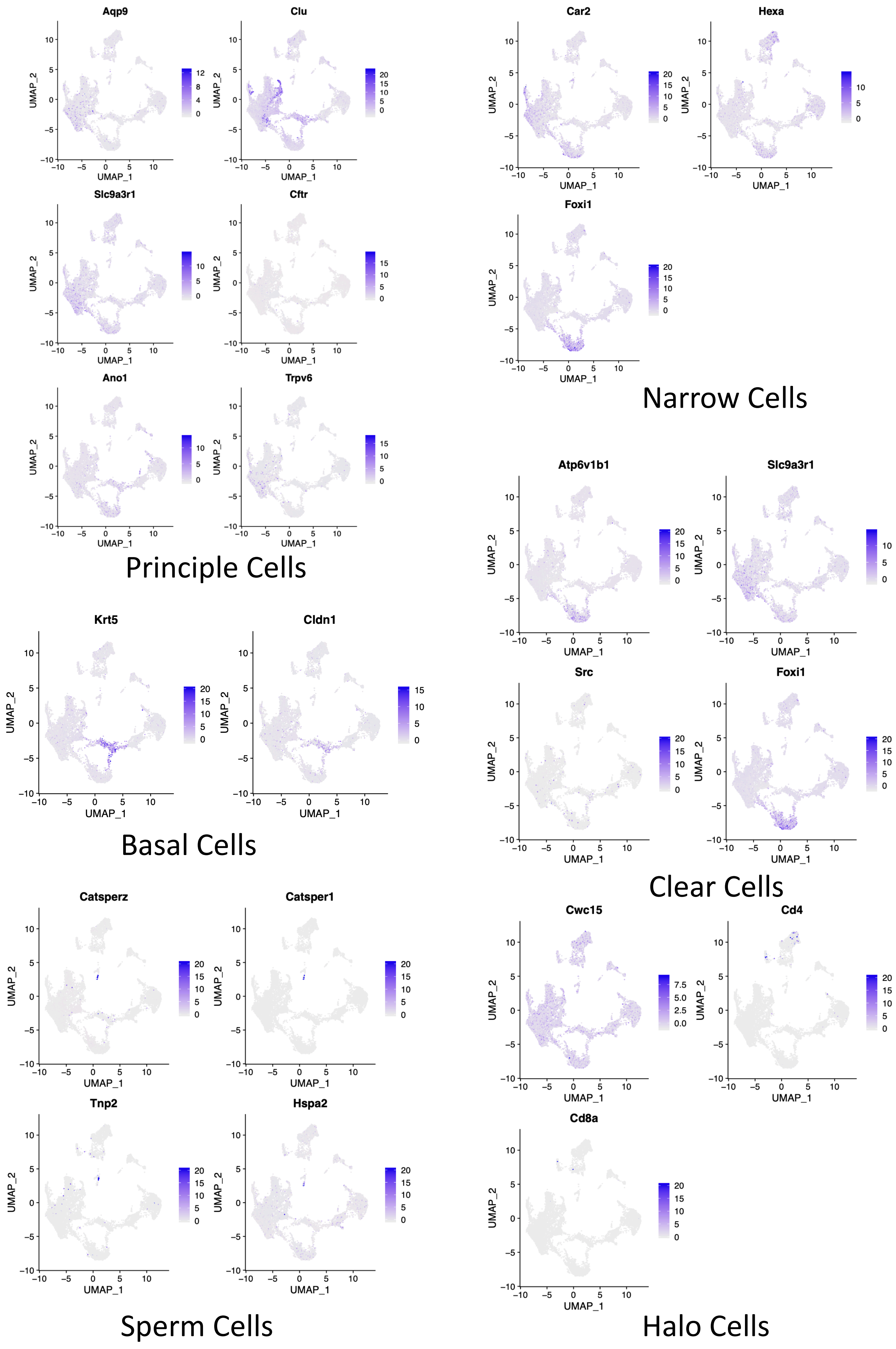

### Figure S2

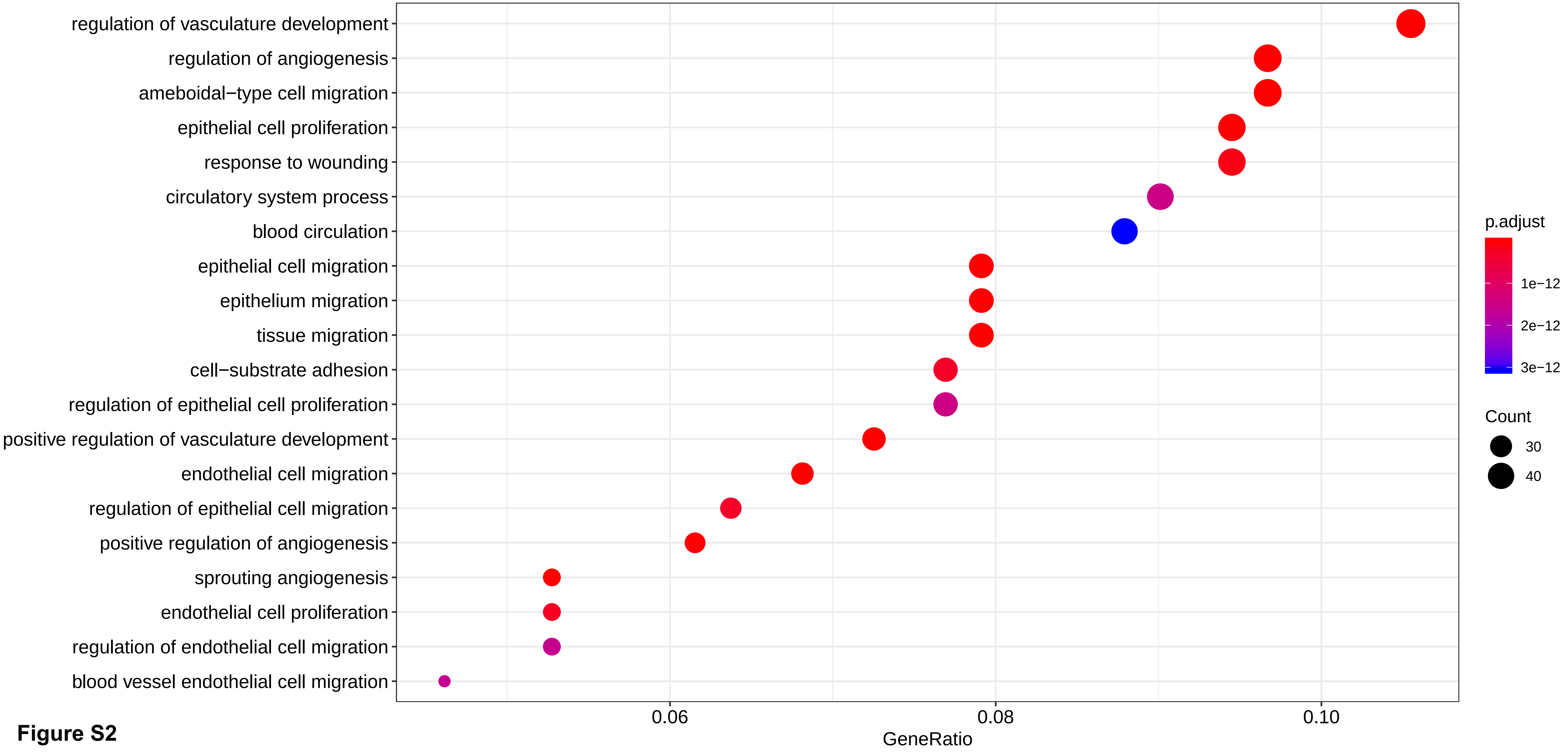

### Figure S3

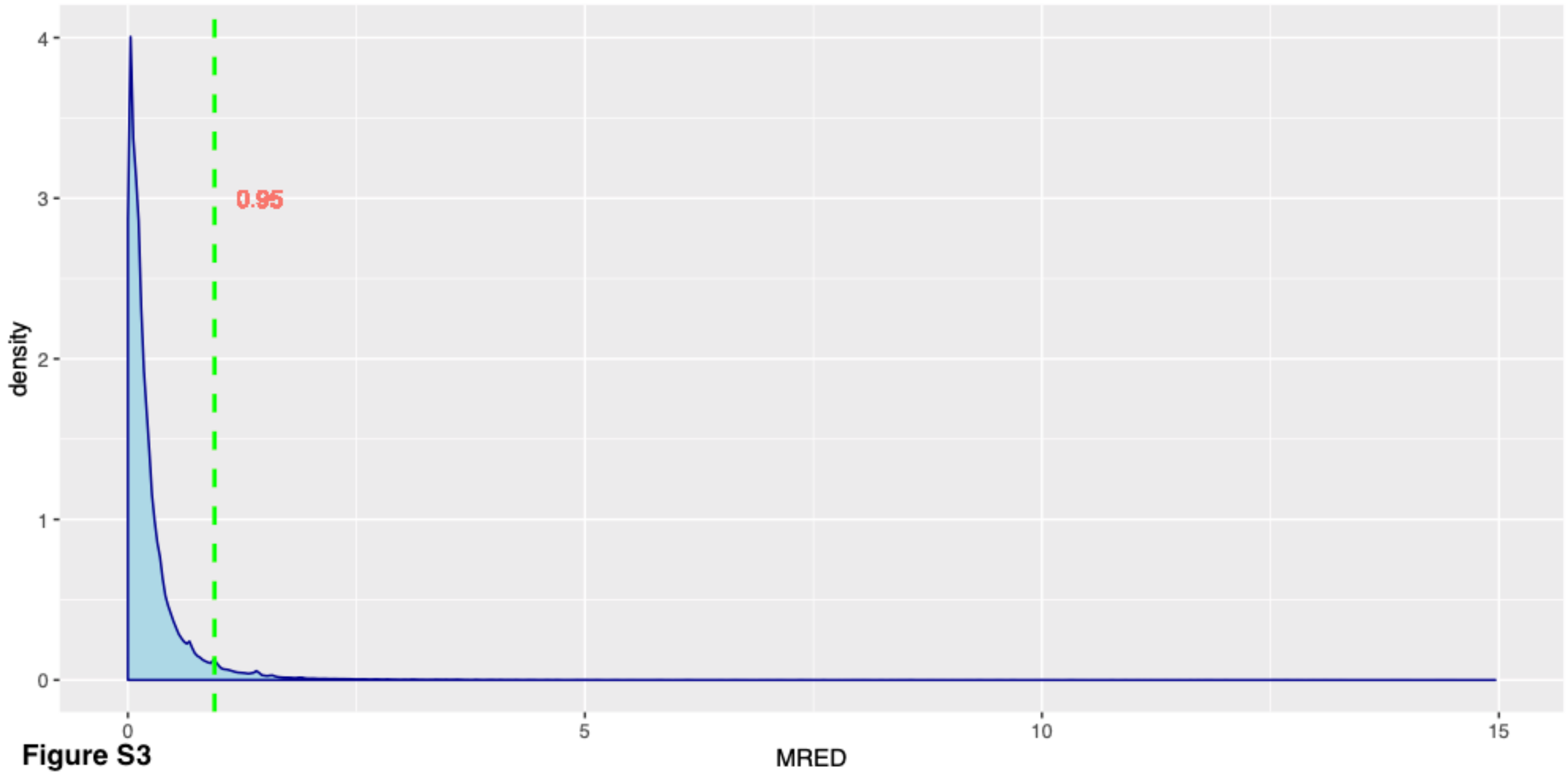


**Figure S3**
